# Neurofibromin deficiency alters the patterning and prioritization of motor behaviors in a state-dependent manner

**DOI:** 10.1101/2024.08.08.607070

**Authors:** Genesis Omana Suarez, Divya S. Kumar, Hannah Brunner, Jalen Emel, Jensen Teel, Anneke Knauss, Valentina Botero, Connor N. Broyles, Aaron Stahl, Salil S. Bidaye, Seth M. Tomchik

## Abstract

Genetic disorders such as neurofibromatosis type 1 increase vulnerability to cognitive and behavioral disorders, such as autism spectrum disorder and attention-deficit/hyperactivity disorder. Neurofibromatosis type 1 results from loss-of-function mutations in the neurofibromin gene and subsequent reduction in the neurofibromin protein (Nf1). While the mechanisms have yet to be fully elucidated, loss of Nf1 may alter neuronal circuit activity leading to changes in behavior and susceptibility to cognitive and behavioral comorbidities. Here we show that mutations decreasing Nf1 expression alter motor behaviors, impacting the patterning, prioritization, and behavioral state dependence in a *Drosophila* model of neurofibromatosis type 1. Loss of Nf1 increases spontaneous grooming in a nonlinear spatial and temporal pattern, differentially increasing grooming of certain body parts, including the abdomen, head, and wings. This increase in grooming could be overridden by hunger in food-deprived foraging animals, demonstrating that the Nf1 effect is plastic and internal state-dependent. Stimulus-evoked grooming patterns were altered as well, with *nf1* mutants exhibiting reductions in wing grooming when coated with dust, suggesting that hierarchical recruitment of grooming command circuits was altered. Yet loss of Nf1 in sensory neurons and/or grooming command neurons did not alter grooming frequency, suggesting that Nf1 affects grooming via higher-order circuit alterations. Changes in grooming coincided with alterations in walking. Flies lacking Nf1 walked with increased forward velocity on a spherical treadmill, yet there was no detectable change in leg kinematics or gait. Thus, loss of Nf1 alters motor function without affecting overall motor coordination, in contrast to other genetic disorders that impair coordination. Overall, these results demonstrate that loss of Nf1 alters the patterning and prioritization of repetitive behaviors, in a state-dependent manner, without affecting motor coordination.

## Introduction

Human genetic disorders can alter brain circuits and behavior, affecting processes ranging from motor function to cognition. The mutations underlying such disorders impact cellular processes such as neuronal growth and differentiation, synaptic transmission and synaptic plasticity, and neuronal excitability [1-3]. These cellular changes cascade through the nervous system, altering complex behaviors. Nervous system function is altered in fragile X syndrome, Rett syndrome, Angelman syndrome, and Williams syndrome, among others, impacting cognition and behavior [4-7]. Given the heterogeneity of neuronal subpopulations – excitatory vs. inhibitory, etc. – as well as the plethora of connectivity architectures across brain regions, a given mutation could produce a range of effects across cells and circuits. Unraveling the complex contributions to behavioral symptoms will require understanding both how genetic mutations alter the function of individual neurons and how they interact at the circuit and systems levels.

Neurofibromatosis type 1 is a genetic disorder that results from loss of function mutations of the NF1 gene that encodes the neurofibromin protein (Nf1). This disorder causes tumor formation and predisposition to cognitive and behavioral symptoms. In addition to its core (mainly cutaneous) symptoms, patients frequently experience comorbidities such as attention-deficit/hyperactivity disorder (ADHD) (∼50%) [8] and autism spectrum disorder (ASD) (∼25%) [9], along with seizures, poor visuospatial skills, executive function deficits, disrupted sleep, repetitive behaviors, and/or pain [8-16]. How Nf1 deficiency alters neuronal activity through effects on interacting circuits across the nervous system is not well understood. Yet alterations to neuronal function likely underlie the increased susceptibility to comorbidities such as ADHD and ASD[8, 14].

Alterations in complex motor function may contribute to the ADHD and ASD symptoms in neurofibromatosis type 1. Such motor functions include grooming and locomotion, which are repetitive, sequenced behaviors that follow hierarchical syntax rules. Animals as diverse as rats and flies groom their body parts in a hierarchical sequence, beginning at the head and proceeding caudally down the body [17-20]. The sequence is dynamically modulated by sensory feedback from each body part [19-21]. Self-grooming is a useful model of behavioral dysregulation in animal models for disorders such as autism spectrum disorder (ASD) – multiple ASD risk factor genes alter grooming when mutated in animal models [18, 22, 23]. Similarly, walking patterns are the result of sequential, coordinated leg movements that modulate forward walking, backward walking, turning, speed changes, and stopping [24]. Children with attention-deficit/hyperactivity disorder (ADHD) exhibit alterations in gait [25], suggesting that disease-driving mutations can affect motor coordination while also driving behavioral symptoms.

*Drosophila* expresses a Nf1 ortholog, and deficiency in this protein leads to behavioral and physiological phenotypes reminiscent of those in mammals. *Drosophila nf1* mutants exhibit learning and memory deficits [26-29], circadian rhythm and sleep disruption [22, 30-32], reduced body size [33], increased grooming [22, 23], impaired jump reflex habituation [34], social (courtship) alterations [35], altered metabolism [31, 32, 36], and tactile hypersensitivity [37]. *Drosophila* neurofibromin contains a conserved catalytic Ras GAP-related domain (GRD) that binds Ras and accelerates its inactivation. Loss of Nf1 in flies increases Ras activity and downstream ERK phosphorylation. In addition, Nf1 deficiency decreases cAMP levels and PKA activity [38-42]. Thus, loss of neurofibromin produces conserved molecular, cellular, and organismal phenotypes through effects on central cellular signaling cascades.

As neurofibromatosis type 1 is associated with a wide range of cognitive and behavioral symptoms, understanding how loss of NF1 function alters repetitive, sequenced behaviors will provide insight into how such genetic disorders impact nervous system function. In this study, we examined how neurofibromin deficiency alters the temporal organization and prioritization of sequenced behaviors, including grooming, locomotion, and feeding. Results revealed that loss of neurofibromin altered grooming and locomotion in a circuit and state-dependent manner.

## Results

### Neurofibromin deficiency drives distinct spatiotemporal grooming patterns across the body

Neurofibromin deficiency increases the time that flies spend grooming [22, 23], potentially reflecting alterations in the activity of command circuits that regulate this motor behavior [19-21, 43]. To dissect the role that neurofibromin plays in the nervous system, we first carried out behavioral analysis of grooming over time. Control (wCS10) flies were compared to *nf1*^*P1*^ mutants, which harbor a genomic deletion encompassing most of the Nf1 locus (including the catalytic GAP-related domain) [33]. We first confirmed that loss of Nf1 increases grooming in both males and females (Figure 1A). Next we examined grooming over time. Individual flies were placed into an open field arena and grooming was quantified over 5-min time windows, starting at t = 0, 30, 60, and 150 min (Figure 1B). When placed into the arena, control flies exhibited an initial period of high grooming (t = 0 min) that decreased by 30 minutes (Figure 1C,E). In contrast, *nf1*^*P1*^ mutants groomed at elevated levels for at least 60 minutes (Figure 1C,F). At 30 and 60 minutes, grooming frequency of the *nf1* mutants was significantly higher than the controls (Figure 1C). By 150 min, grooming in both control and *nf1* mutants had decreased and were no longer significantly different from one another (Figure 1C). Depending on the time, median grooming in the *nf1* mutants increased 72% - 403%. To investigate the neuronal contributions to this effect, we knocked down neurofibromin with RNAi using the Gal4/UAS system. RNAi expression was targeted to neurons using the pan-neuronal Gal4 driver R57C10-Gal4 [44]. Upon introducing these flies to the open field arena (t = 0 min), there was significantly higher total spontaneous grooming in the experimental group compared to the heterozygous driver (Gal4/+) and effector (UAS/+) genetic controls (Figure 1D). Flies with pan-neuronal Nf1 knockdown displayed elevated grooming levels for 60 minutes. This was followed by a decrease at 150 minutes (when grooming was no longer significantly different from both controls) (Figure 1D). These results indicate that loss of Nf1 increased spontaneous grooming for at least one hour due to neuronal effects.

**Figure 1.**
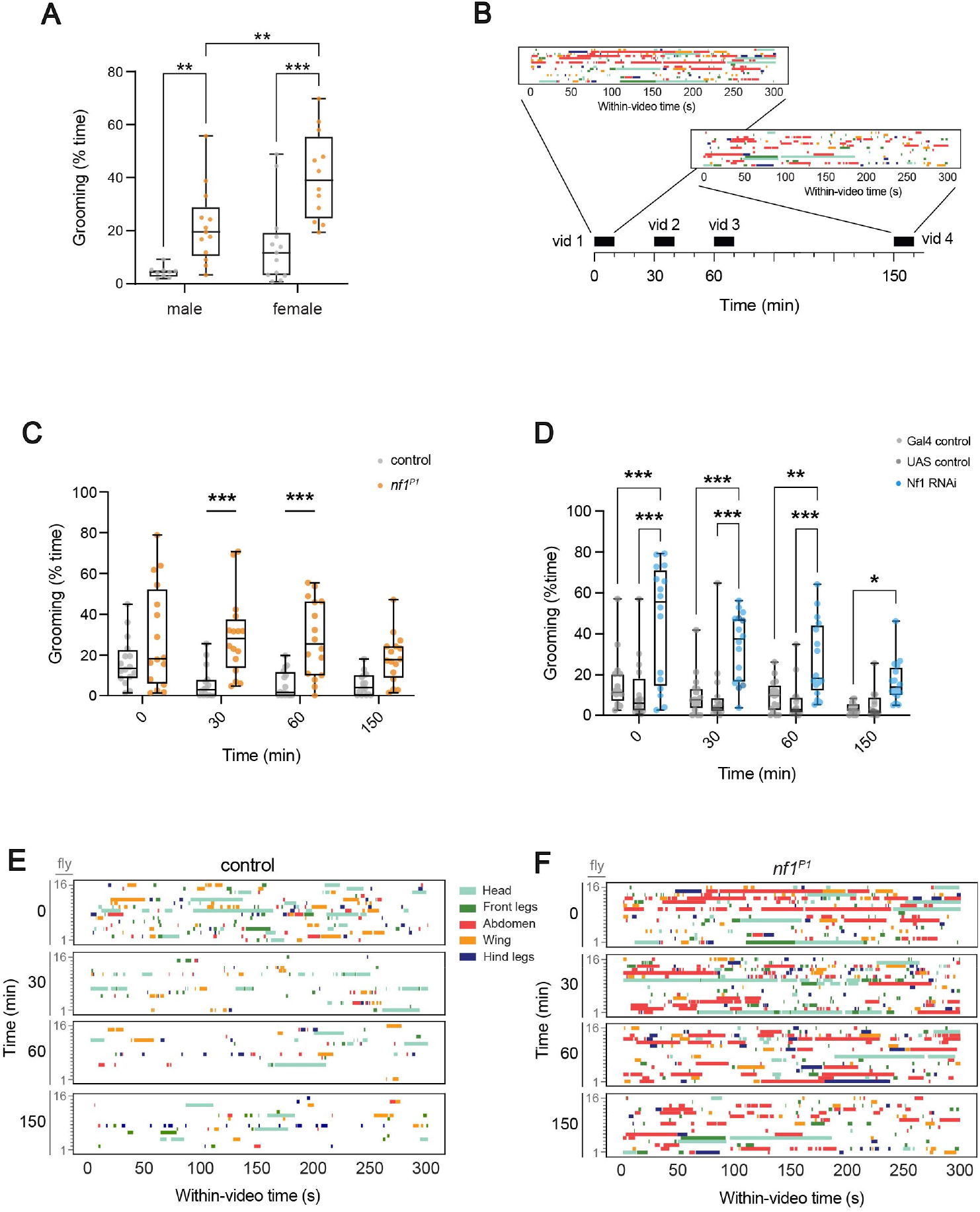
Nf1 deficiency alters grooming frequency across time. Box plots: median = line, box = interquartile range; whiskers = min/max values, individual data points: circles. *p < 0.05, **p < 0.01, ***p < 0.001 (Šidák; n = 12-16). **(A)** Loss of Nf1 increased grooming in both males and females. **(B)** Time course of video collection and example of data (5-min grooming ethograms, replicated from panel F) visualized at two different time points. **(C)** Grooming frequency for control (wCS10) flies and *nf1*^*P1*^ mutants. **(D)** Grooming frequency with pan-neuronal Nf1 knockdown (R57C10-Gal4>UAS-Nf1^RNAi^,UAS-dcr2). **(E)** Ethograms of grooming for control (wCS10) flies, showing each grooming bout across animals, with the groomed body part color-coded. **(F)** Ethograms of grooming for *nf1*^*P1*^ mutants.

Examination of the grooming patterns across body parts revealed heterogeneity in the effects of Nf1 deficiency. Loss of Nf1 exerts behavioral effects via actions on distributed neuronal circuits [23], raising the question of how homogenous or heterogeneous the neuronal effects of Nf1 deficiency are across neuronal circuit elements. To approach this, we investigated how Nf1 deficiency affects the temporal evolution of grooming across different body parts. *nf1*^*P1*^ mutants exhibited significantly increased grooming of the abdomen at 0, 30, and 60 minutes following introduction to the arena, with a trend still present at 150 minutes (Figure 2A,C). Additionally, the back legs exhibited a small Nf1-dependent increase at 60 minutes compared to the control (Figure S1). Grooming of other body parts was not statistically significant from controls (Figure 2A, S1, S2). These findings suggested that the loss of Nf1 via genomic mutation increases grooming frequency, but this increase was nonuniform across body parts.

**Figure 2.**
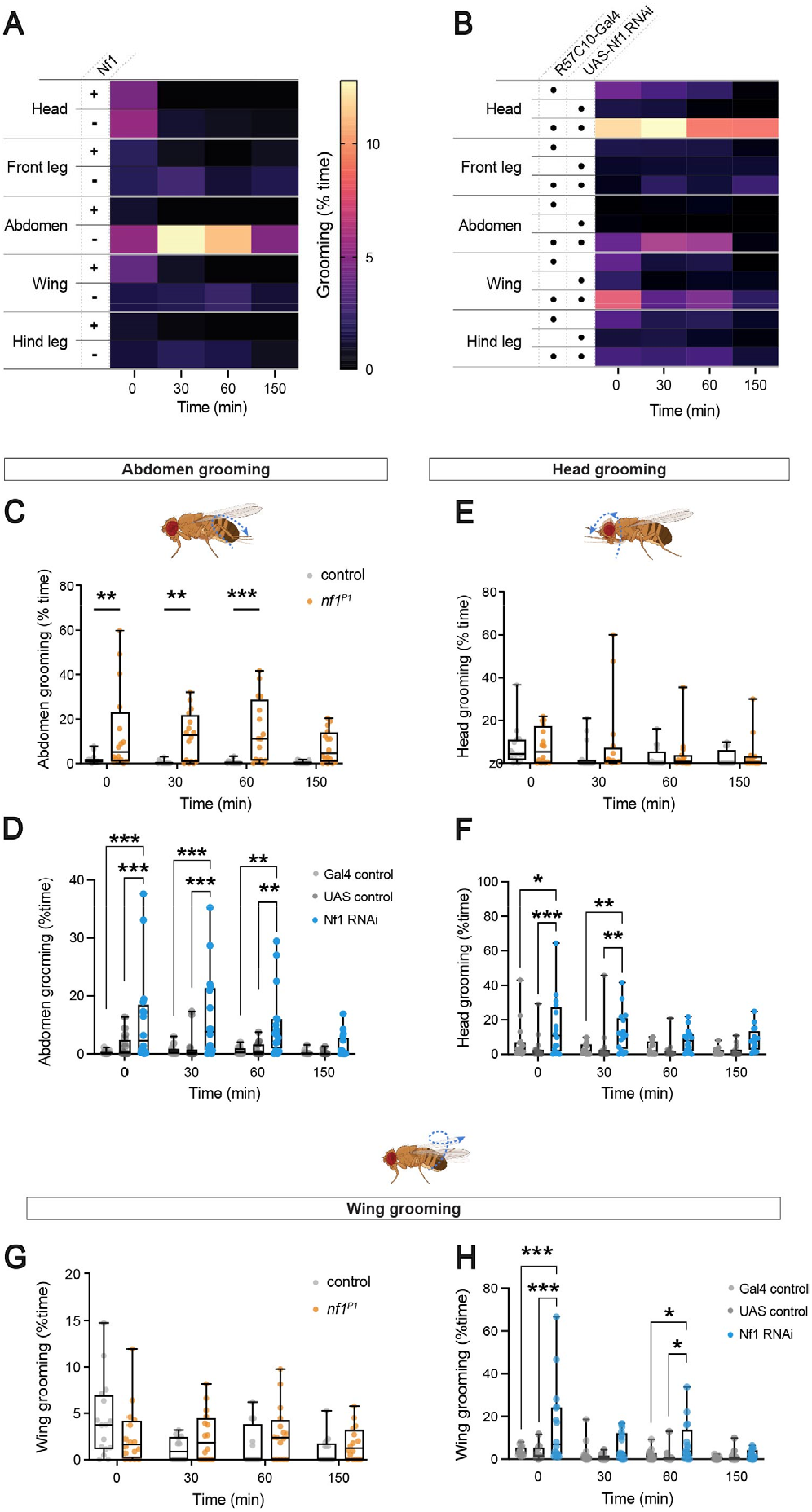
Nf1 deficiency alters grooming in a body part-specific manner. **(A)** Heat map of grooming time across body parts and time in controls and *nf1*^*P1*^ mutants. **(B)** Heat map of grooming time across body parts and time with pan-neuronal Nf1 knockdown (R57C10-Gal4>UAS-Nf1^RNAi^,UAS-dcr2). **(C)** Abdomen grooming across time in controls (wCS10) and *nf1*^*P1*^ mutants. Box plots: median = line, box = interquartile range; whiskers = min/max values, individual data points: circles. **(D)** Abdomen grooming with pan-neuronal Nf1 knockdown (R57C10-Gal4> UAS-Nf1^RNAi^,UAS-dcr2). **(E)** Head grooming in controls and *nf1*^*P1*^ mutants. **(F)** Head grooming with pan-neuronal Nf1 knockdown. **(G)** Wing grooming in controls and *nf1*^*P1*^ mutants. **(H)** Wing grooming with panneuronal Nf1 knockdown. Fly drawing modified from biorender.com.

To further test the heterogeneity of Nf1 effects across grooming circuits, we analyzed the effect of pan-neuronal Nf1 knockdown on grooming. Upon introducing the flies to the open field arena, grooming in control flies was initially elevated and decreased within 30 minutes (Figures 2B,F, S2). In the experimental knockdown group, there was a significant increase in head grooming compared to the Gal4/+ and UAS/+ controls (Figures 2B,F, S2). This persisted for 30 minutes before gradually decreasing. Similarly, there were increases in abdomen and wing grooming (Figure 2B,D,H) that persisted for 30-60 minutes. There were no consistent changes in front or back leg grooming (Figures S1, S2). Overall, grooming increased with loss of Nf1; both *nf1*^*P1*^ mutants and pan-neuronal Nf1 knockdown increased abdomen grooming, though pan-neuronal RNAi also elevated head and wing grooming (Figures 1E,F, 2). These data revealed heterogeneity of Nf1 effects on grooming behaviors across different body parts over time. Differences between the Nf1 mutants and RNAi suggest that loss of Nf1 alters the balance of excitation (and/or inhibition) across neuronal circuits, with different loss of function manipulations producing somewhat different effects.

Changes in total grooming time result from changes in either frequency of grooming initiation and/or duration of grooming bouts. To determine which of these underlies the neurofibromin effect, we analyzed individual grooming bouts, quantifying the bout count and bout duration in *nf1* mutants and controls. The increase in abdomen grooming arose from increases in bout count, with increased bout duration at some time points (Figure S3). There were also increases in front and back leg grooming bout counts at one time point (30 min) and an increase in back leg grooming bout duration at one time point (60 min) (Figure S4). Taken together, increases in both bout count and bout duration contribute to the increase in total grooming time (with the bout count being the more robust). These data suggest that loss of neurofibromin enhances the activation of grooming circuits, resulting in more frequent grooming initiation and increased perseveration (i.e., sustained grooming).

### Neurofibromin deficiency modulates a state-dependent switch from grooming-to foraging-dominant behavioral modes

As Nf1 deficiency increases grooming, grooming command circuits are aberrantly engaged. To test how this behavior is prioritized in the face of competing stimuli, we examined the hunger state dependence of spontaneous grooming. When food is withheld for some time, flies increase their locomotion to forage for food, a phenomenon known as starvation-induced hyperactivity [45-47]. When they are walking, they cannot be grooming. Thus, we wondered whether the reduction in grooming over time in the open field (Figure 1C) (which lacks food) could be attributed to altered hunger state. To test this, we first compared flies in arenas without food to those with ad libitum food access throughout the experiment (Figure 3A). In the presence of food, *nf1*^*P1*^ mutants displayed significantly higher grooming compared to controls at all time points, including the longest time point (150 min) (Figure 3A). Thus, *nf1* mutant flies continue to groom significantly more than controls if food is available.

**Figure 3:**
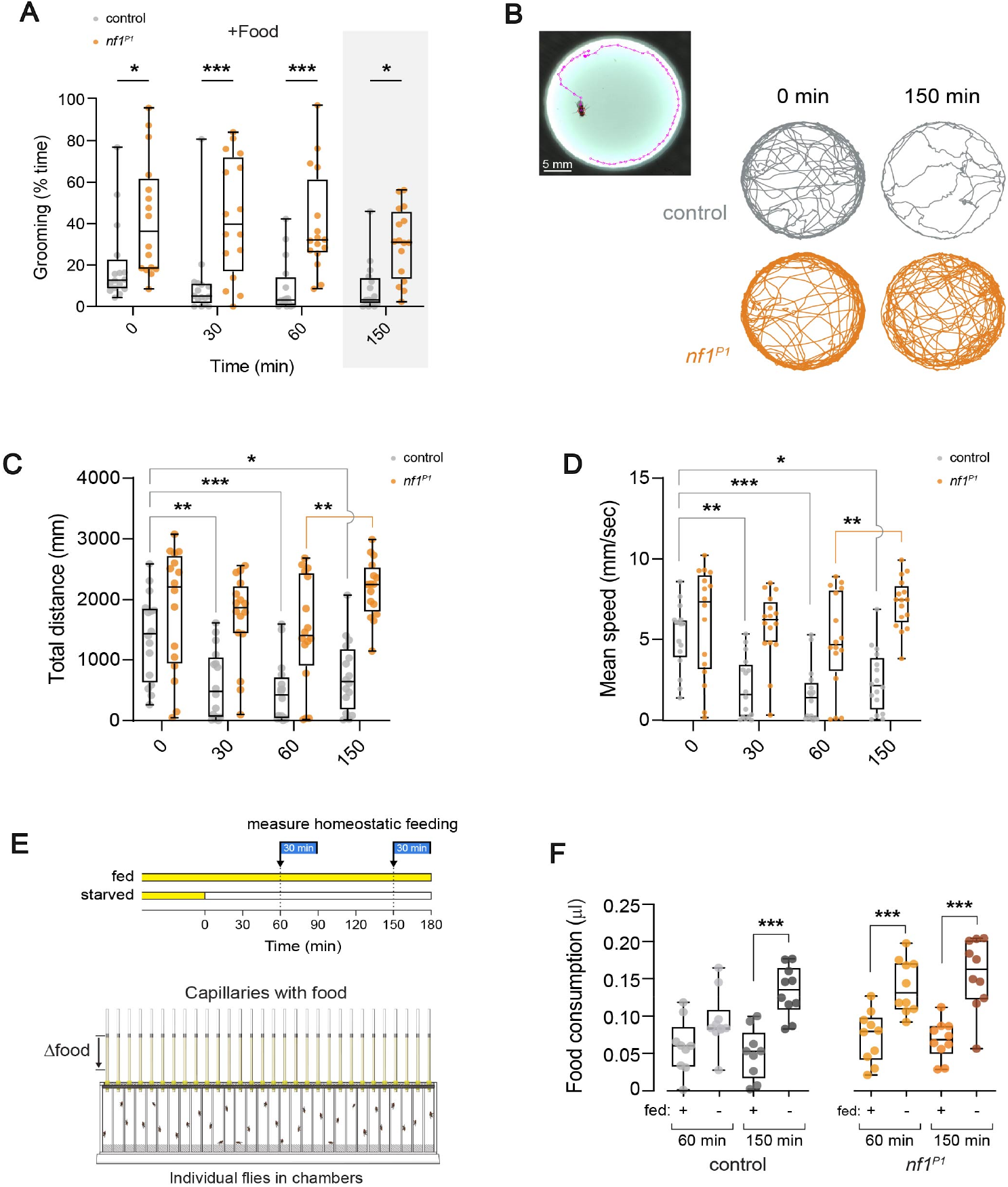
Nf1 deficiency modulated a state-dependent behavioral switch from grooming to locomotion. **(A)** Quantification of grooming (% time) in an open field arena when solid food was provided ad libitum (+Food), comparing control (wCS10) flies to *nf1*^*P1*^ mutants. **(B)** Still capture of fly locomotion tracking, with xy position tracks over 5 min for a representative control fly and *nf1*^*P1*^ mutant at 0 and 150 min **(C)** Total distance traveled for control flies and *nf1*^*P1*^ mutants. **(D)** Mean walking speed for control flies and *nf1*^*P1*^ mutants. **(E)** Diagram of the capillary feeding assay and protocol to test homeostatic feeding. Two cohorts each of controls and *nf1*^*P1*^ were tested: one kept on food continuously (“fed”) and one in which food was withheld at starting at t = 0 (“starved”). Food consumption was measured starting at t = 0 and t = 150 minutes. **(F)** Feeding in controls and *nf1*^*P1*^ mutants, comparing feeding in fed (fed +) and starved (fed -) conditions.

This suggests that when food is absent, grooming may decrease in a hunger state-dependent manner.

To investigate whether the decrease in grooming frequency at longer time points could be driven by foraging, we quantified locomotion. Locomotion was tracked over time, analyzing distance traveled and walking speed. In open field arenas, flies exhibit an initial stage of high walking activity that gradually decreases over time (considered to be exploratory behavior) [48, 49]. In control flies, we observed this as higher locomotor activity upon introduction to the arena, which dropped within 30 minutes (Figure 3B,C). This was reflected in both a significant decrease in the distance traveled and speed from the 0 to 30 min time points (Figure 3 C,D). While the median locomotor activity decreased in *nf1*^*P1*^ mutants across these time points, the effect did not reach statistical significance (Figures 3C,D, S5). However, from 60 to 150 minutes, a significant increase in locomotion was observed in the *nf1*^*P1*^ mutants for both distance and speed (Figures 3C,D, S5). These data are consistent with the interpretation that the reduction in grooming frequency results from an increase in locomotion, providing additional evidence that neurofibromin deficiency prompts a behavioral shift toward foraging behavior.

Grooming decreased over time in the open field, while locomotion increased, raising the question of whether hunger state drives the behavioral changes. To determine whether hunger is a cue was a driver of increased locomotion, we examined food intake for signs homeostatic feeding, characterized by a rebound in feeding driven by negative energy balance [50, 51]. Food intake was measured after periods of food deprivation using a capillary feeding assay [50]. Two cohorts of flies were tested for each genotype (control and *nf1*^*P1*^): one group that had continuous ad libitum access to food (“fed”) and one in which food was withheld for either 60 or 150 minutes (“starved”) (Figure 3E). These time points corresponded to the duration of food deprivation experienced in the open field experiments. In control flies, we did not observe a significant difference in the amount of liquid food consumed between the fed and starved groups at the 60-minute time point (Figure 3F). However, at 150 minutes, starved control flies consumed significantly more food compared to the fed controls, indicating increased hunger (Figure 3F). In the case of *nf1*^*P1*^ mutants, homeostatic feeding was observed at both time points (Figure 3F), demonstrating that food deprivation increases hunger faster in neurofibromin-deficient flies. This increase in hunger corresponds to a higher metabolic rate in *nf1*^*P1*^ mutants [31], and could drive starvation-induced hyperactivity along with decreased grooming.

### Loss of Nf1 in sensory and grooming command neurons modulates grooming in a circuit-specific manner

Grooming of each body part is controlled by discrete sensory neurons and grooming command neurons [19, 20, 43, 52, 53], raising the question of whether loss of Nf1 affects the sensory neurons and/or command neurons directly. To test this, we employed a targeted approach involving the knockdown of Nf1 in selected neuronal subsets [23]. We considered a range of plausible circuit architectures interconnecting the command circuits. Nf1 could affect the sensory input to the command neurons, the command neurons themselves, interactions between the sensory and command neurons (Figure 4A), or via interconnected neurons across more broad swaths of the central nervous system. Based on these hypothetical network architectures, we generated four testable models: 1) sensory input modulation, 2) command circuit modulation, 3) sensory + command circuit modulation, and 4) systems-level modulation. These models were challenged with a series of circuit-specific Nf1 manipulations.

**Figure 4:**
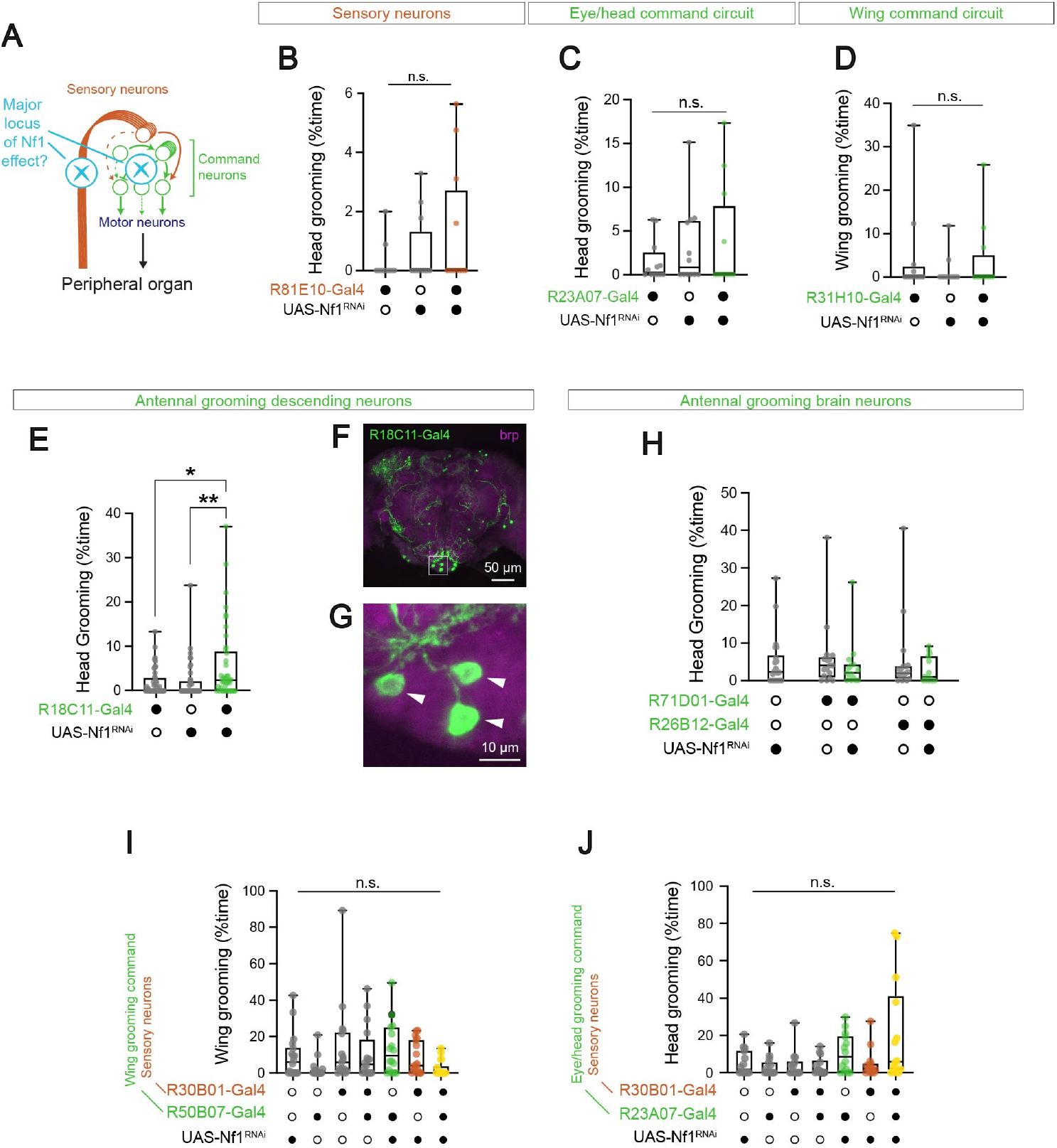
Knocking down Nf1 in sensory neurons and/or grooming command circuits shifted the pattern of grooming without affecting total grooming time. Box plots: median = line, box = interquartile range; whiskers = min/max values, individual data points: circles. *p < 0.05, **p < 0.01, n.s. = not significant (Šidák; n = 16). **(A)** Simplified diagram of sensory neurons and antennal grooming command neurons, potential sites of modulation by Nf1 deficiency. RNAi was targeted to sensory neurons, command neurons, or both. **(B)** Effect of Nf1 knockdown in sensory neurons on head grooming (R81E10-Gal4>UAS-Nf1^RNAi^,UAS-dcr2). Experimental flies were compared to heterozygous Gal4/+ and UAS/+ controls. **(C)** Effect of Nf1 knockdown in eye/head grooming command neurons on head grooming (R23A07-Gal4>UAS-Nf1^RNAi^, UAS-dcr2). **(D)** Effect of Nf1 knockdown in wing grooming command neurons on wing grooming (R31H10-Gal4>UAS-Nf1^RNAi^, UAS-dcr2). **(E)** Effect of Nf1 knockdown in antennal grooming command neurons on head grooming (R18C11-Gal4>UAS-Nf1^RNAi^, UAS-dcr2). **(F)** Expression pattern of neurons labeled by R18C11-Gal4, focusing on the central brain. Box highlights the inset shown in panel G. GFP:green; brp:magenta. **(G)** Expanded view of the boxed region from panel F, including the somata of antennal descending neurons (white arrowheads). **(H)** Effect of Nf1 knockdown in antennal descending neurons (aDN) on head grooming using two different drivers (R71A06-Gal4 or R26B12-Gal4). **(I)** Effects of Nf1 knockdown in sensory neurons (R30B01-Gal4), wing grooming command neurons (R50B07-Gal4), and both, on wing grooming. **(J)** Effects of Nf1 knockdown in sensory neurons (R30B01-Gal4), eye/head command neurons (R23A07-Gal4), and both, on head grooming.

To test whether loss of Nf1 in sensory neurons alone increased grooming frequency, we knocked Nf1 down in sensory neurons with RNAi using two Gal4 drivers (R81E10 and R30B01-Gal4) [21, 44]. These drivers label multiple subsets of sensory neurons including mechanosensory bristles, the Johnston’s organ, chemosensory receptors, chordotonal organs, and campaniform sensillae [21]. There was no significant effect on grooming frequency when Nf1 was knocked down in the sensory neurons labeled by either of these drivers (Figures 4 B,I,J, S6). Overall, these data suggested that Nf1 deficiency in sensory neurons did not account for the increased grooming when Nf1 was reduced across the nervous system. Next, we tested the impact of Nf1 knockdown in command circuits on grooming frequency and temporal structure, utilizing six different Gal4 drivers (R23A07, R18C11, R31H10, R71D01, R26B12, and R50B07-Gal4) that encompass grooming command neurons. These drivers include command neurons for circuits that control grooming of the eye/head (R23A07), antennae (R18C11, R71D01 [aBN1], and R26B12 [aBN2]), and wings (R31H10 and R50B07) [19-21, 19]. No significant changes in overall grooming frequency were observed when Nf1 was knocked down using any these drivers (Figures 4 C,D,E,H; S6). Examining grooming of each body part individually, we found that knocking down Nf1 in antennal descending neurons (via R18C11) modestly biased grooming toward the head (Figure 4E). This driver labels three pairs of antennal descending neurons that modulate antennal grooming (Figure 4F,G) [19]. The behavioral effect of Nf1 knockdown in these neurons suggests that loss of Nf1 in one component of the neuronal circuits driving grooming may affect grooming patterning. In contrast, knocking down Nf1 in aBN1 and aBN2 antennal grooming command neurons did not detectably alter head grooming (Figure 4H). Thus, the large increase in grooming with broad knockdown likely representing either additive effects across multiple components of the grooming command circuits and/or their inputs.

To test whether Nf1 functions additively in pairs of neuron types within the grooming circuitry, we knocked Nf1 down in both command neurons and/or and their sensory inputs. This was done using one Gal4 driver and/or in grooming command neurons with a second Gal4 driver. Two combinations sensory+command drivers were R30B01+R50B07-Gal4 of neuron tested: and R30B01+R23A07-Gal4. The R30B01-Gal4 driver provides broad coverage of sensory neurons innervating mechanosensory bristles across the eye, head, abdomen, wing, notum, and leg, as well as sensory neurons innervating the Johnston’s organ, chemosensory receptors, chordotonal organs, and campaniform sensillae [21]. Thus, the R30B01+R50B07-Gal4 combination includes both mechanosensory inputs to the grooming command circuits (R30B01) and the wing grooming command circuit (R50B07). Similarly, the R30B01+R23A07 covers mechanosensory neurons and an eye/head grooming command circuit. Neither of these two driver pairs significantly altered grooming frequency when used to knock down Nf1 (Figure 4H-J). These data suggest that Nf1 deficiency in sensory circuits does not interact additively with command neurons, and that it is required more broadly and/or in higher-level neurons.

### Loss of Nf1 alters the pattern and prioritization of stimulus-evoked grooming behaviors

Loss of neurofibromin increased the frequency of spontaneous grooming, but its effect on stimulus-evoked grooming is unknown. When flies are covered with a fine layer of dust, they vigorously groom to remove the dust [20, 54]. This stimulus-evoked grooming follows a temporal progression (approximately cephalocaudal) resulting from hierarchical command circuit recruitment [20]. To test whether neurofibromin alters the pattern and prioritization of sensory-evoked grooming movements, we dusted flies and quantified dust removal. Within 35 minutes of dusting, controls removed much of the dust from their head, thorax, wings, and abdomen (Figure 5). *nf1*^*P1*^ mutants similarly removed most of the dust from their head, thorax, and abdomen, but wing cleaning was significantly reduced – i.e., the dust was removed from the wings more slowly in mutants. This represented a departure from the normal cephalocaudal grooming sequence, wing grooming shifted down in priority. Loss of Nf1 affected spontaneous grooming and stimulus-evoked grooming in different ways. Spontaneous grooming was elevated in the abdomen, head, and wings (Figure 2). Yet the deficit in stimulus-evoked grooming was focused on the wings. Overall, these data suggest that both the patterning and prioritization of stimulus-evoked grooming was altered by loss of neurofibromin. Further, since different body parts were affected in different conditions, loss of Nf1 affects multiple circuits, potentially including those upstream of the grooming command neurons.

**Figure 5:**
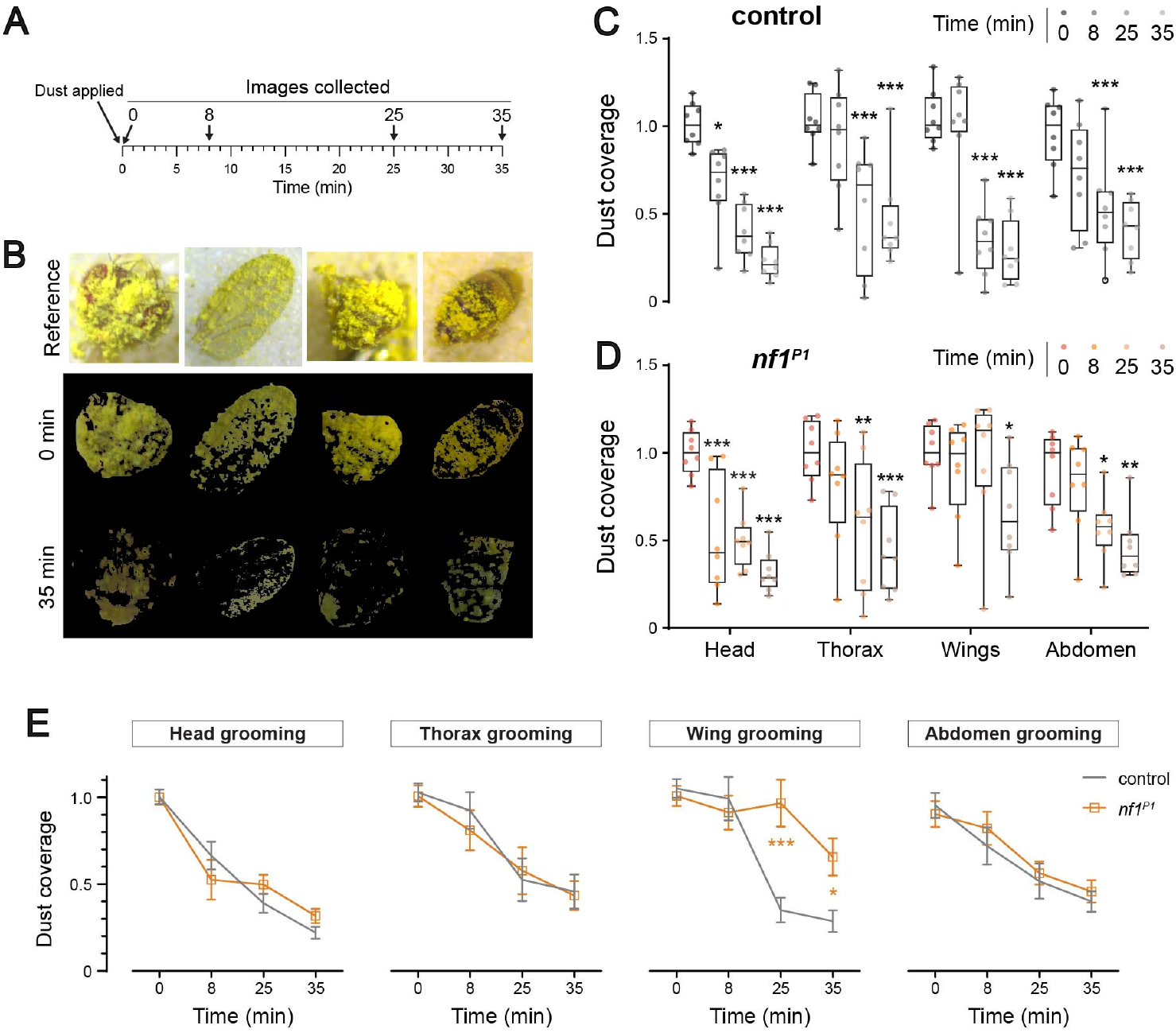
Loss of Nf1 altered the temporal evolution of stimulus-evoked grooming. **(A)**. Diagram of the experimental protocol. Flies were dusted, and amount of dust remining on each body part was imaged at t = 0, 8, 25, and 35 min. **(B)**. Reference images of different body parts immediately after dusting, with images showing dust coverage after 0, 8, 25, and 35 min. **(C)**. Dust removal in control (wCS10) flies. Dust coverage is the fraction of dust relative to those imaged at time 0. *p < 0.05; **p < 0.01, ***p < 0.001 re: time 0 (ANOVA/Šidák, n = 8). **(D)**. Dust removal in *nf1*^*P1*^ mutants, plotted as in panel C. **(E)**. Time course of dust removal (same data as in panels C,D) comparing controls and *nf1*^*P1*^ mutants at each time point. *p < 0.05; **p < 0.01, ***p < 0.001, comparing controls and *nf1*^*P1*^ mutants (ANOVA/Šidák, n = 8). Error bars = S.E.M.

### Nf1 deficiency alters forward walking velocity without any major defects in leg kinematics

Given that Nf1 deficient flies show altered locomotion, we wondered whether their walking gait was impaired (a common feature of genetic disorders affecting motor function). To test this, we used a 3D leg kinematics analysis pipeline [55] to compare the fine structure of leg kinematics during walking in controls and flies with genomic Nf1 deletion (*nf1*^*P1*^). For this analysis, flies were tethered to a pin and allowed to walk on a spherical treadmill (Figure 6A). Nf1 mutants exhibited increased walking speed in the tethered preparation compared to K33 (w+) controls (Figure 6B), similar to untethered flies in the open field (Figure 3B-D). Pan-neuronal knockdown of Nf1 produced a similar effect (Figure 6B). Despite the change in walking speed, overall gait was not altered compared to controls. Leg placement was similar in *nf1* mutants and control flies (Figure 6C,D). Stance, step, and swing durations were also similar between genotypes across the instantaneous velocity range (Figure 6E-G). This suggests that flies lacking Nf1 increased their walking speed in the same way as control flies – by changing the frequency of stepping (decreasing stance duration and increasing stance length) (Figure 6E-G and [56]). Further, inter-leg coordination was normal in mutants – there was no major difference in coupling of leg movements between either legs that move in phase (e.g., left front leg and right middle) (Figure 6K,L) or antiphase (e.g., left and right front legs) (Figure 6I,J) during tripod gait. Taken together, these results indicate that Nf1 deficiency altered walking speed without affecting gait. Loss of Nf1 is unlikely to affect lower-level motor control areas like proprioceptors and motor neurons, as alterations in those would lead to major anomalies in leg kinematics. Therefore, Nf1 likely impacts locomotion via higher-order walking control centers.

**Figure 6:**
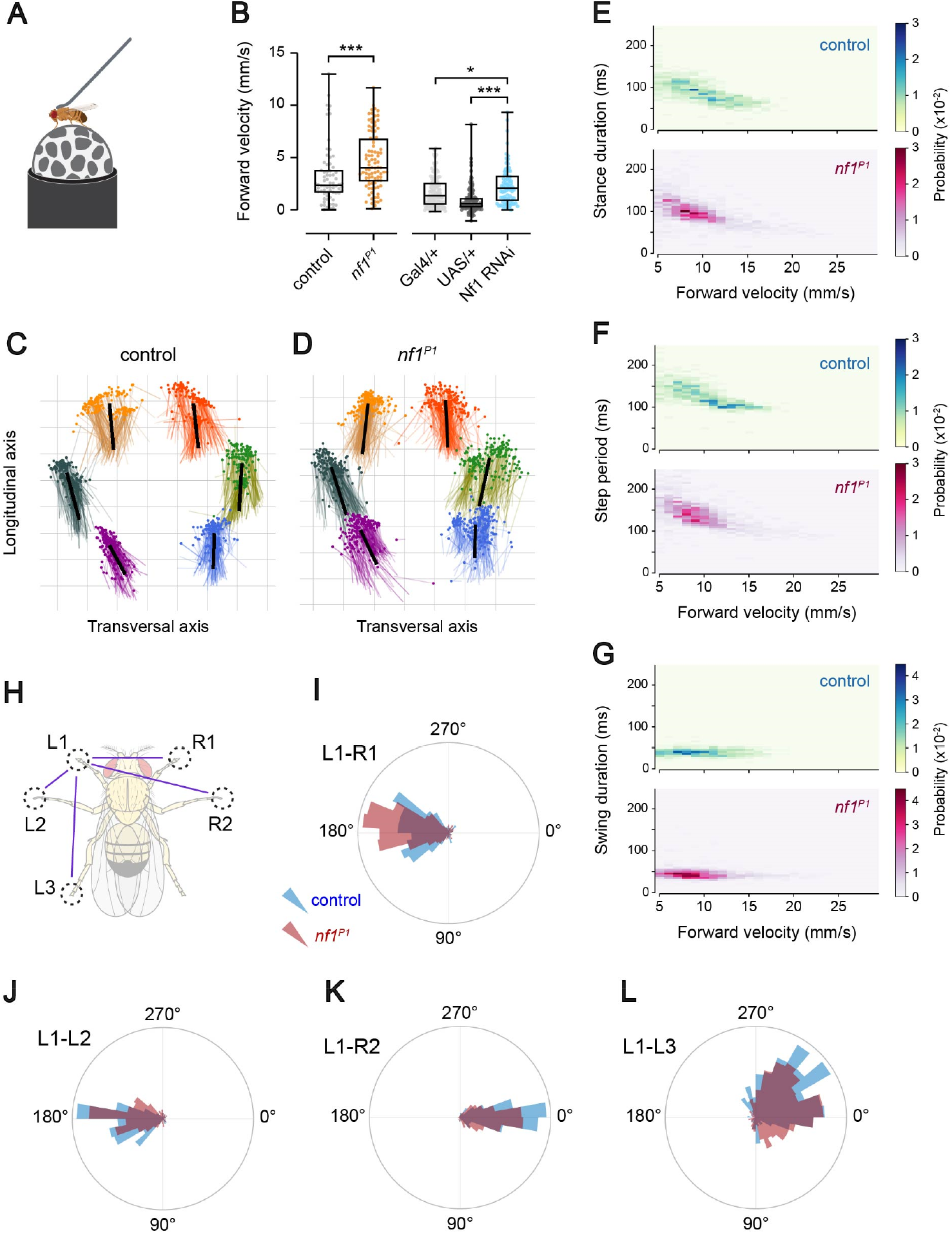
Nf1 deficiency increased walking speed without altering gait or kinematics. **(A)**. Diagram of the experimental setup. Fly drawing modified from biorender.com. **(B)**. Forward walking speed of controls (K33) vs. *nf1*^*P1*^ mutants (***p < 0.001 [Mann-Whitney], n = 70-90) and with pan-neuronal Nf1 knockdown (R57C10-Gal4>UAS-Nf1-RNAi) (*p < 0.05, ***p < 0.001 [Kruskal-Wallis/Dunn], n = 80-124). **(C)**. Stance trajectory in control flies. 300 individual points plotted (randomly selected from >1000 steps). The dots indicate touch down locations, which are connected to the liftoff with a line. The black line is the mean of all trajectories. **(D)**. Stance trajectory in *nf1*^*P1*^ mutants, plotted as in panel C. **(E)**. Stance duration for the L1 leg, comparing K33 controls and *nf1*^*P1*^ mutants. Probability density is graphed as a heat map. **(F)**. Step period for the L1 leg. **(G)**. Swing duration for the L1 leg. **(H)**. Diagram of leg coordination phase comparisons shown in panels I-L. **(I)**. L1-R1 leg movement phase plot, comparing K33 controls and *nf1*^*P1*^ mutants. **(J)**. L1-L2 leg movement phase plot. **(K)**. L1-R2 leg movement phase plot. **(L)**. L1-L3 leg movement phase plot.

## Discussion

The present data demonstrate that loss of Nf1 alters motor pattern structure and prioritization of motor behaviors in *Drosophila*. To ensure survival, animals must prioritize one behavior and inhibit another based on internal and external cues [57]. We found that Nf1 is required to maintain normal activity levels across multiple behaviors over time, as Nf1-deficient flies exhibit altered grooming and walking. The grooming can be overridden by hunger, which shifts the balance toward walking to prioritize foraging. These findings suggest that loss of Nf1 alters neuronal activity, increasing activation of spontaneous grooming and walking speed, as well as changing the prioritization of stimulus-evoked grooming. Importantly, motor coordination is not notably altered, and grooming remains plastic and state dependent.

Grooming is a structured and sequenced set of motor patterns that are modulated by sensory inputs and prioritized according to body part [20]. In both flies and mammals, grooming follows an approximately anterior to posterior sequence. In flies, grooming of each body part is controlled by discrete command circuits. Each command circuit comprises multiple elements: sensory neurons that provide the input to command neurons that initiate grooming and send output to descending neurons to execute the motor patterns [19-21, 43, 53]. Hierarchical regulation ensures that, when sensory stimuli drive the need to clean multiple body parts, grooming occurs in a prioritized sequence, with certain body parts (such as the eyes and the antennae) taking precedence over others (such as the wings and thorax) [21].

Loss of Nf1 increases spontaneous grooming frequency [22], likely through effects on distributed circuits [23]. This created two likely a priori scenarios regarding the impact of Nf1 on grooming; loss of Nf1 could 1) increase the grooming across all body parts, or 2) increase the grooming of body parts at the pinnacle of the grooming hierarchy, which would then be expected to suppress the grooming of other body parts via inhibitory circuit effects. However, our data revealed a third, unexpected outcome: Nf1 genomic mutation predominantly increased spontaneous grooming of the abdomen (with knockdown also increasing head and wing grooming). This suggests that the effects of loss of Nf1 could be heterogenous across different command circuits/neurons, leading to distinct alterations in grooming patterns for different body parts. The pattern of grooming alterations differed between the *nf1* genomic mutation and pan-neuronal knockdown with RNAi. There are several potential reasons for this. The expression level of Nf1 could be important, with functional differences between reduction (RNAi) and elimination (genomic mutant) of Nf1. Secondly, variations in the expression level of the R57C10-Gal4 driver (or efficacy of the RNA-induced silencing complex) across different sets of neurons may drive distinct grooming patterns. Third, the expression pattern of Gal4 lines can shift over the course of development [58], potentially introducing heterogeneity of Nf1 knockdown during critical periods [23]. Finally, non-neuronal cells could contribute to the observed phenotype; while R57C10 drives relatively selective Gal4 expression in neurons [44], there are some scattered non-neuronal cells in its expression pattern [59]. In a slightly different arena/environment, head grooming was favored (rather than abdomen) with loss of Nf1 [23], suggesting that context may influence the pattern as well.

Increased grooming is due to loss of Nf1 in excitatory cholinergic circuits [23]. Relatedly, Nf1 deficiency causes alterations in synaptic transmission at the larval neuromuscular junction that originate, at least in part, from central excitatory circuits [37]. If the effects of Nf1 deficiency are selective for excitatory neurons, then the number and connectivity of excitatory vs. inhibitory neurons in each grooming circuit would influence how loss of Nf1 affects grooming of each body part. Future studies are needed to address the heterogeneity of cell-autonomous effects of Nf1 deficiency, as well as the contributions of variations in neuronal architecture. Alterations in excitatory:inhibitory (EI) balance are hypothesized to contribute to cognitive and behavioral alterations across of range of genetic disorders [60]. Given the position of Nf1 as modulator of central neuronal signaling pathways including Ras, cAMP, and G protein-coupled receptor signaling, it is plausible that EI balance is altered in ways that affect cognition and behavior.

Loss of Nf1 dramatically elevates grooming (72% - 403%), yet this grooming remains plastic and can be overridden by stimuli that are prioritized over grooming (e.g., foraging). In open field arenas without food, the flies became hungry and accumulated detectable negative energy balance after 60-150 minutes (as assayed by homeostatic feeding). Increasing locomotion is a foraging strategy to locate food under starvation conditions [45-47]. By 150 minutes in the open field without food, grooming frequency dropped and locomotion increased, reflecting a state-dependent switch from a grooming-to foraging-dominant behavioral mode. Thus, grooming circuits are engaged by Nf1 deficiency until hunger increases and overrides the grooming drive. The increase in hunger preceded the increase in locomotor activity, suggesting that flies accumulate negative energy balance for a period before foraging is engaged. This is likely a strategy to maximize energy utilization, as increasing locomotor activity under energy-restricted conditions accelerates starvation; it represents a high-risk, last-resort strategy to find food. Overall, this suggests that loss of Nf1 alters network dynamics, reshaping the activation and perseveration of specific motor behaviors in a state-dependent way, but sparing some behavioral plasticity.

Nf1 mutants exhibited increased walking speed on a spherical treadmill, yet exhibited normal leg kinematics and gait patterns. This suggests that Nf1 affects the macro properties of locomotion like walking speed without causing defects in micro-properties like inter-leg coordination or stepping direction. Other mutations that impact motor function, such as Parkinson’s disease (PD) (α-synuclein expression, parkin mutants) and spinocerebellar ataxia (SCA) models (mutant SCA3 expression) [61] affect locomotion differently. For instance, PD and SCA disease models exhibit markedly altered gait, generating uncoordinated movements and erratic foot placement [62]. Thus, Nf1 is unlikely to affect low-level motor control neurons (central pattern generators, proprioceptors, and descending neurons). Rather it may act on higher-order walking control neurons [55, 56, 63], directly and/or indirectly (e.g., via metabolic changes [31]).

Nf1 exerts effects on neuronal function during a critical period of development [23]. Specifically, there is a critical period for Nf1 effects on grooming in *Drosophila* spanning the late 3rd instar larval phase and first half of the pupal phase. This developmental period encompasses a series of neurodevelopmental steps, including cell proliferation, migration, differentiation, dendritic remodeling, axon guidance, synaptogenesis, and survival [64-68]. Therefore, Nf1 effects on the neuronal circuitry regulating grooming could result from alterations in one or more of these processes. Ultimately, the behavioral effects likely emanate from altered adult neuronal composition, connectivity, excitability, or synaptic transmission. Some Nf1 mutations alter neuronal excitability in rodents [1]. Although this is associated with tumor formation, it suggests that changes in neuronal excitability may be a conserved mechanism. At the molecular level, the behavioral effect of Nf1 requires an intact Ras GTPase-activating domain, suggesting that altered Ras signaling could underlie (or contribute to) the phenotype. Ras influences multiple developmental processes, including playing roles in growth, migration, cytoskeletal integrity, survival, and differentiation [69]. Though many of the described effects of Nf1 deficiency involve Ras signaling [2, 31, 39, 70], loss of Nf1 also alters cAMP signaling, which contributes to neuronal phenotypes [1, 26, 27, 71, 72]. Therefore, cAMP signaling could contribute to grooming phenotypes, potentially downstream of Ras [42]. Future studies will be necessary to delineate the specific contributions of each pathway, as well as dissect the key downstream signaling molecules and interactions.

Overall, the present data reveal that loss of Nf1 affects the neuronal network activity regulating temporally-sequenced behaviors in a circuit-specific and state-dependent manner. Changes in grooming were nonuniform across body parts, suggesting heterogeneity of effects on the command circuits that drive grooming of each body part. Loss of Nf1 exerted strong behavioral effects, but those could be overridden by stimuli that are prioritized for survival. In addition, the activation of locomotor behavior was altered, with changes in walking speed (increased forward walking speed on a spherical treadmill), but no detectable change in kinematics or gait. Thus, the loss of Nf1 alters the frequency, patterning, prioritization, and execution of sequenced motor behaviors. These data lay the groundwork for future studies to address the molecular mechanisms of Nf1 effects on neurons, circuit alterations, and the prioritization and sequencing of behaviors following mutation of genes that drive neurodevelopmental disorders.

## Acknowledgements

We thank Timothy Stelmat for designing and constructing lighted enclosures for the open field experiments. Flies obtained from the Bloomington *Drosophila* Stock Center (NIH P40OD018537) and the Vienna *Drosophila* Resource Center (VDRC, www.vdrc.at) were used in this study. Imaging was carried out at the University of Iowa Central Microscopy Research Facility (CMRF). Acquisition of the CMRF Leica SP8 Laser Scanning Confocal microscope with STED capability was made possible by a generous grant from the Roy J. Carver Charitable Trust; additional CMRF funding was provided by the University of Iowa Office of the Vice President for Research, the Carver College of Medicine, and the College of Liberal Arts and Sciences. This research was supported by NIH/NINDS R01 NS097237, R01 NS126361, R01 NS114403, and R21 NS124198. We thank Lisa Ringen, Linda Buckner, Rob Svetly, Kathleen O’Brien, and Melissa Benilous for administrative support. Formatting of this preprint is based on the bioRxiv template by Christian L. Ebbesen (https://github.com/chrelli).

## Materials and Methods

### Drosophila maintenance

Flies were raised on cornmeal/agar food medium, housed in incubators at 25 °C, 60% relative humidity, and kept on a 12:12 light:dark cycle according to standard protocols. Behavioral assays were performed using 3-8 days old flies. Except where indicated, males were used (to prevent egg accumulation). The *nf1*^*P1*^ mutation was backcrossed 6 generations into the wCS10 genetic background. The Nf1 RNAi line was obtained from the Vienna *Drosophila* RNAi Center (VDRC #109637); UAS-dicer2 was coexpressed to potentiate the RNAi effect44 and was included with all experimental and UAS/+ genotypes. The empty attP control line (VDRC #60100) was used in Gal4/+ control crosses to match the genetic background across groups. The following lines used in this study were obtained from the Bloomington *Drosophila* Stock Center (BDSC): R81E10-Gal4 (BDSC #48367), R52A06-Gal4 (38810), R23A07-Gal4 (49010), R18C11-Gal4 (48808), R31H10-Gal4(48104), R30B01-Gal4 (49517), and R50B07-Gal4 (38729).

### Grooming analysis

Grooming was quantified as previously described [22, 23]. Flies were placed into an open field arena, 25.4 mm in diameter and 2.85 mm in height, consisting of an opaque (white) PLA lateral boundary covered on the top and bottom with two clear polycarbonate sheets. The apparatus was illuminated from below with white light emitting diodes that were filtered through a sheet of white acrylic; light intensity was measured at 720 lm/m2 in the location of the fly. A camera (FLIR Teledyne Blackfly S) fitted with a 25mm lens (Edmund Optics) was mounted above each arena. A single male fly was placed in the arena with an aspirator and recorded at 7.5 frames per second, 1616 × 1240 with lossless Motion JPEG 2000 compression. 5-min videos were recorded at several intervals, starting at: 0, 30, 60, and 150 min following introduction into the open field arena. Grooming was manually scored via frame-by-frame analysis, recording the start and stop frames for each grooming event. Grooming events were categorized according to body part: front legs, head/eye, abdomen, wings, or hind legs. The percentage of time spent grooming, bout count, and bout duration was calculated. Grooming was graphed in aggregate or by individual body parts, as appropriate. Ethograms were generated with a custom Python script and heat maps were plotted using GraphPad Prism. Stimulus-evoked grooming was performed as previously described [20]. Each fly was coated in Reactive Yellow 86 dust and transferred to an open field area with a mesh floor to allow dust to fall through. Flies were allowed to clean for a period of time, then anesthetized with CO2 and the head, thorax, abdomen, and wings were separated and photographed individually. Using the color selection tool in Adobe Photoshop, yellow pixels were selected. and then manually corrected as needed to ensure that dust particles were accurately selected. Yellow pixels were quantified and the number divided by total number of pixels to calculate the percentage of dust coverage on each body part. The percent coverage for each body part/fly/time point was normalized to the median value of the dust coverage at time 0.

### Open field locomotion tracking

Locomotion was tracked (in videos collected as above) using MATLAB with DLTdv8 [73]. Accuracy of the tracking was visually confirmed and any errors in the xy position tracking were manually corrected. Total distance and mean walking speed were calculated over the duration of the video from the frame-by-frame xy positions.

### High-resolution 3D leg kinematics analysis

Locomotor gait and kinematics were examined using a tethered fly preparation as previously described [55, 74-76]. 7-10 day old male flies were tethered to a 34-gauge needle with ultraviolet light-cured glue and placed on a spherical treadmill (6mm diameter) suspended in a stream of compressed air. The flies were placed on the ball with minimum wait time after tethering and allowed up to 5 min of recovery before starting the experiment. The compressed air was passed through an in-line heating element to bring the local temperature on the ball up to 32 °C, inducing high-speed spontaneous walking. Rotational velocity of the ball was monitored in real-time using two orthogonally placed motion sensors at 50Hz. Each trial was triggered in a closed loop fashion when the forward velocity crossed a threshold empirically determined to signify sustained walking. Each fly could trigger a maximum of 10 trials, but flies that triggered more than 5 trials were included in the final dataset for analysis. Each trial was 7s long. Eight cameras (FLIR BFS-U3-16S2M-CS) fitted with InfiniStix 194100 lenses and near-infrared bandpass filters (Midopt BP850) were placed surrounding the ball so that all legs were continuously visible from at least one pair of cameras. The fly was illuminated with a custom infrared ring emitting focused light to the plane of the ball. The cameras, infrared light source and the ball tracker were all synchronized and triggered at 200 Hz by an Arduino microcontroller, with camera exposure time set to 200μs. The fly was recorded with a resolution of 1440×1072 pixels. Each fly was left on the ball for a maximum of 20 min.

### Capillary feeding assay

Feeding was quantified in adult male flies using a capillary feeding assay [50]. Individual flies were placed into chambers (46 mm × 7 mm) containing 1% agar at the bottom and a glass capillary at the top. The glass capillaries contained an aqueous food solution (5% sucrose + 5% yeast extract) and dye to track food consumption. One group of 10 flies was fed for 150 min, one group of 10 flies was starved for 60 minutes, and one group of 10 flies was starved for 150 minutes prior to providing food for the experiment. Total feeding was measured in a 30-min window and analyzed in Fiji.

### Immunohistochemistry

Five to seven-day-old adult brains were dissected in 1% paraformaldehyde in S2 medium and processed as previously described [77]. Samples were stained with primary antibodies for 3 hours at room temperature and at 4 °C overnight, followed by secondary antibodies for 3 hours at room temperature and 5 days at 4 °C. Incubations were performed in 3% normal goat blocking serum. Samples were mounted in Vectashield (Vector Laboratories) for analysis. The following antibodies were used: rabbit anti-GFP (1:1000, Invitrogen), mouse anti-nc82 (1:50, DSHB), goat anti-rabbit IgG, and goat anti-mouse IgG (1:800, Alexa 488 or Alexa 633, respectively, Invitrogen). Images were obtained using a Leica SP8 Laser Scanning Confocal microscope with LAS X software.

### Statistical analysis

Normality of data was assessed with the Shapiro-Wilk Normality Test. In figures, box plots graph the median as a line, the interquartile range (IQR) as a box, and whiskers extend to the min/max values. Hypothesis testing was carried out using the Wilcoxon rank-sum test or Kruskal-Wallis omnibus test followed by Dunn multiple comparisons tests. Two-way comparisons were carried out with a two-way ANOVA followed by Šidák’s multiple comparisons tests. For RNAi analysis, the experimental group was compared to both heterozygous genetic controls and considered positive if it significantly differed from both controls in the same direction. Statistics and graphing were carried out with Graphpad Prism, version 10.1.1.

