## Supplemental Figures for "Neurofibromin deficiency alters the patterning and prioritization of motor behaviors in a state-dependent manner"

### *Supplementary Figure Legends*

**Figure S1**. Nf1 deficiency produces little effect on leg grooming. Box plots: median = line, box = interquartile range; whiskers = min/max values, individual data points: circles. *p < 0.05, ***p < 0.001 (Šidák; n = 16). **(A)** Front leg grooming in controls (wCS10) and in $nf1^{P1}$ mutants across time. **(B)** Front leg grooming over time with pan-neuronal Nf1 knockdown (R57C10-Gal4>UAS-$Nf1^{RNAi}$, UAS-dcr2). **(C)** Back leg grooming in control and $nf1^{P1}$ flies across time. **(D)** Back leg grooming over time with pan-neuronal Nf1 knockdown.

**Figure S2**. Effects of Nf1 deficiency across all body parts and time points. Data are from the same experiments graphed in Figures 2 and S1. *p < 0.05, **p < 0.01, ***p < 0.001 (Šidák; n = 16). **(A)** Grooming across body parts and time in control (wCS10) flies vs. $nf1^{P1}$ mutants. **(B)** Grooming across body parts and time with pan-neuronal Nf1 knockdown (R57C10-Gal4>UAS-$Nf1^{RNAi}$,UAS-dcr2).

**Figure S3**. Nf1 deficiency altered abdomen grooming initiation and duration, increasing bout count and bout duration. Box plots: median = line, box = interquartile range; whiskers = min/max values, individual data points: circles. *p < 0.05, ***p < 0.001 (Šidák; n = 16). **(A)** Head grooming bout count across time in control (wCS10) flies and $nf1^{P1}$ mutants. **(B)** Head grooming bout duration across time in controls and $nf1^{P1}$ mutants. **(C)** Abdomen grooming bout count across time in controls and $nf1^{P1}$ mutants. (D) Abdomen grooming bout duration across time in controls and $nf1^{P1}$ mutants. **(E)** Wing grooming bout count across time in controls and $nf1^{P1}$ mutants. **(F)** Wing grooming bout duration across time in controls and $nf1^{P1}$ mutants.

**Figure S4**. Nf1 deficiency altered leg grooming initiation (bout count) and duration (bout duration) at certain time points. Box plots: median = line, box = interquartile range; whiskers = min/max values, individual data points: circles. *p < 0.05, **p < 0.01, ***p < 0.001 (Šidák; n = 16). **(A)** Front leg grooming bout count across time in control (wCS10) flies and $nf1^{P1}$ mutants. **(B)** Front leg grooming bout duration across time in controls and $nf1^{P1}$ mutants. **(C)** Back leg grooming bout count across time in controls and $nf1^{P1}$ mutants. **(D)** Back leg grooming bout duration across time in controls and $nf1^{P1}$ mutants.

**Figure S5**. Locomotion in representative controls and $nf1^{P1}$ mutants over time. **(A)** xy position tracks for a control (wCS10) fly (same as in Figure 4B) over five-minute videos at 0 min, 30 min, 60 min, and 150 min following introduction to the open-field arena **(B)** xy position tracks for an $nf1^{P1}$ mutant (same as in Figure 4B). **(C)** Grooming ethograms (top) and locomotor activity traces for a control fly at 0 min, 30 min, 60 min, and 150 min following introduction to the open field arena. **(D)** Grooming ethograms (top) and locomotor activity traces for an $nf1^{P1}$ mutant.

**Figure S6**. Knocking down Nf1 in sensory neurons, command neurons, and sensory+command neuron combinations does not affect grooming frequency (total grooming time). Box plots: median = line, box = interquartile range; whiskers = min/max values, individual data points: circles. *p < 0.05, **p < 0.01, n.s. = not significant (Šidák; n = 16). **(A)** Effect of Nf1 knockdown in sensory neurons on grooming time (R81E10-Gal4> UAS-$Nf1^{RNAi}$,UAS-dcr2). **(B)** Effect of Nf1 knockdown in eye/head grooming command neurons on grooming time (R23A07-Gal4>UAS-$Nf1^{RNAi}$, UAS-dcr2). **(C)** Effect of Nf1 knockdown in antennal grooming command neurons on grooming time (R18C11-Gal4>UAS-$Nf1^{RNAi}$, UAS-dcr2). **(D)** Effect of Nf1 knockdown in wing grooming command neurons on grooming time (R31H10-Gal4>UAS-$Nf1^{RNAi}$, UAS-dcr2). **(E)** Effects of Nf1 knockdown in sensory neurons (R30B01-Gal4), wing grooming command neurons (R50B07-Gal4), and both, on grooming time. **(F)** Effects of Nf1 knockdown in sensory neurons (R30B01-Gal4), eye/head command neurons (R23A07-Gal4), and both, on grooming time.

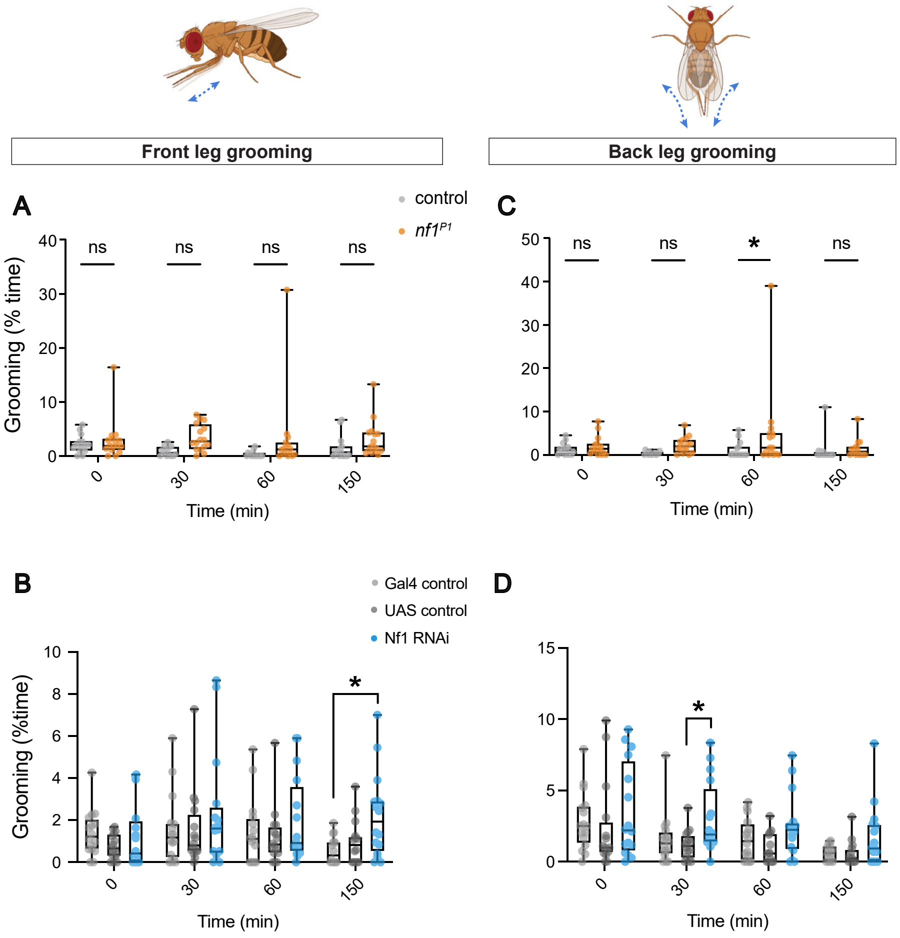
Front leg grooming
Back leg grooming
A
C
B
D
control
nf1P1
ns
*
Grooming (% time)
Grooming (%time)
Time (min)
Gal4 control
UAS control
Nf1 RNAi

Figure S1

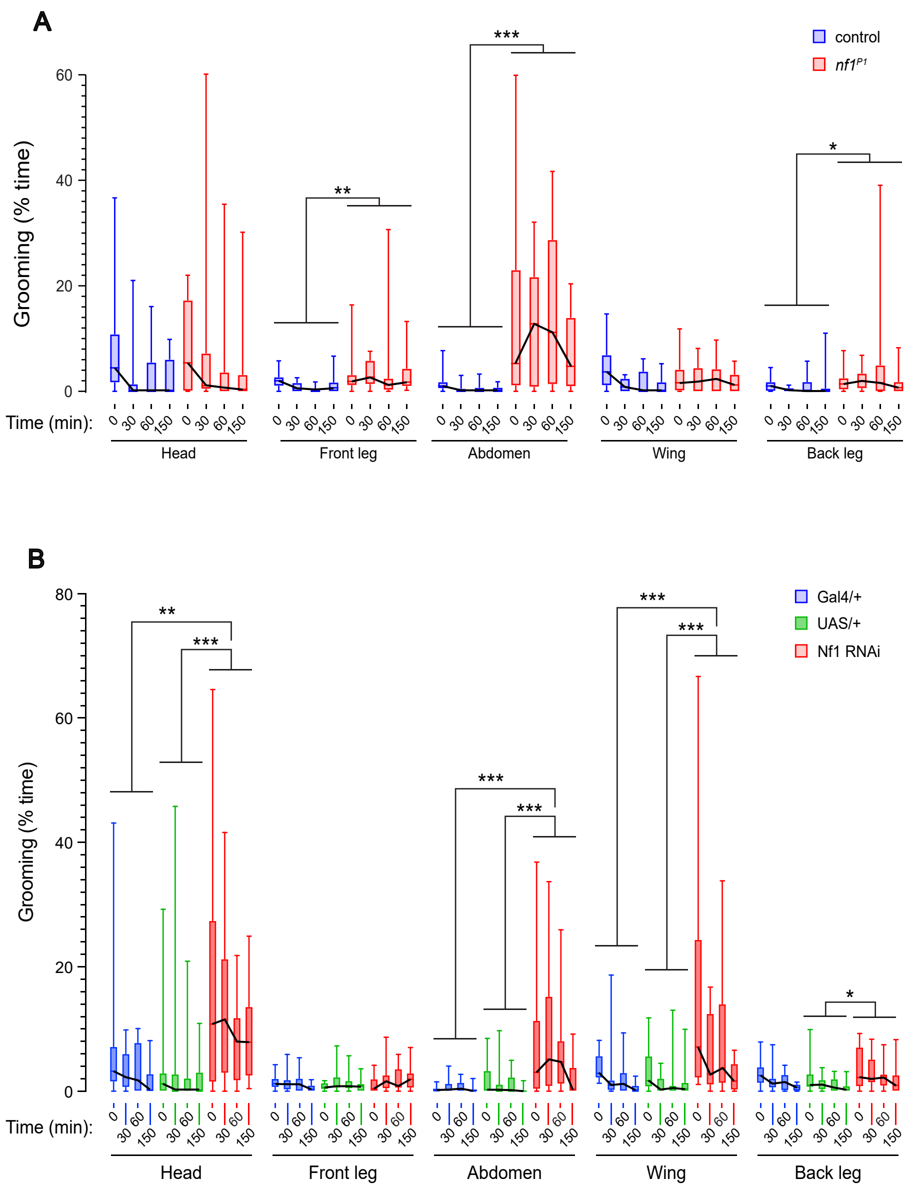
A
control
nf1^P1
Grooming (% time)
Time (min):
Head
Front leg
Abdomen
Wing
Back leg
B
Gal4/+
UAS/+
Nf1 RNAi
Grooming (% time)
Time (min):
Head
Front leg
Abdomen
Wing
Back leg

Figure S2

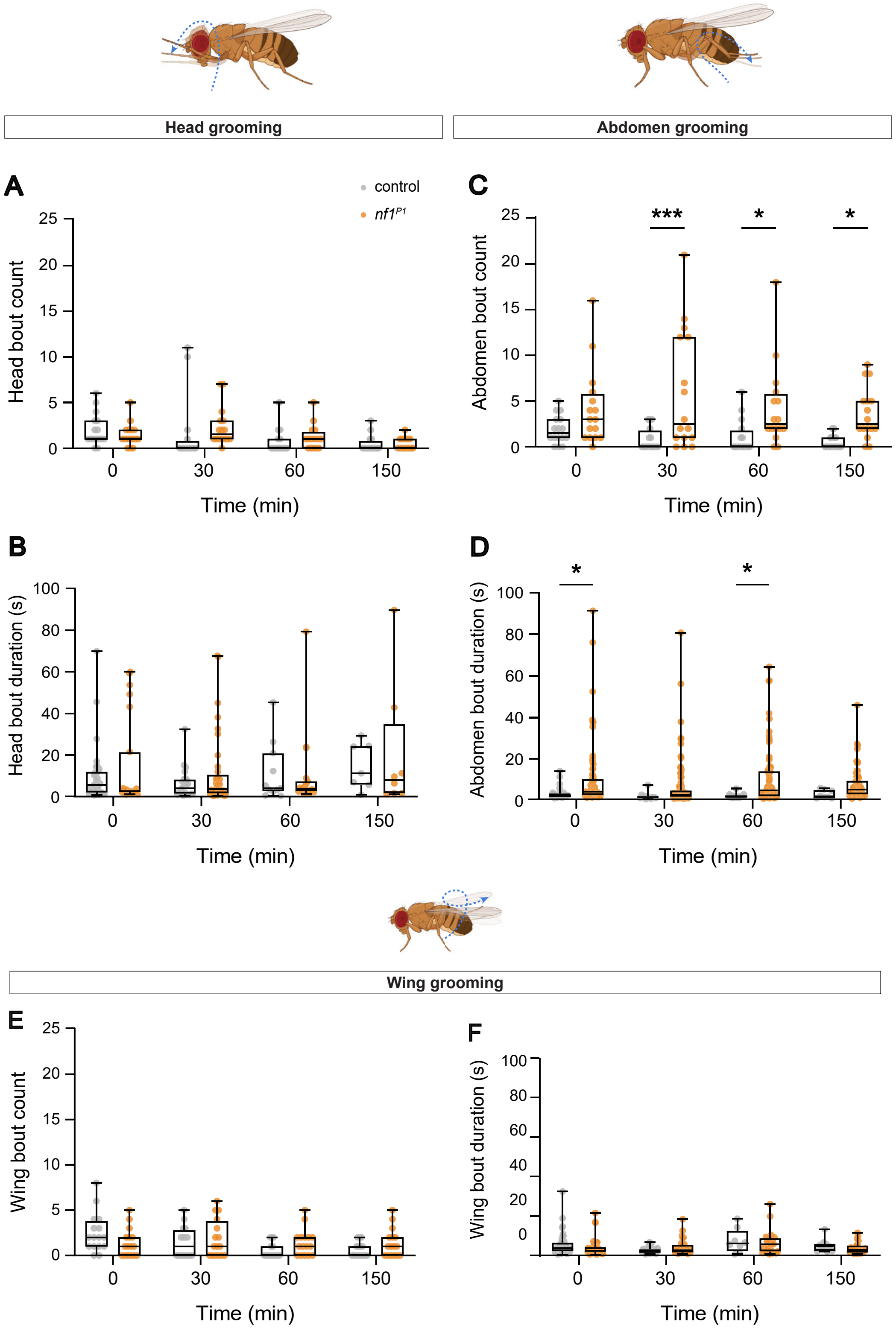
Head grooming
Abdomen grooming
A
control
nf1P1
Head bout count
Time (min)
B
Head bout duration (s)
Time (min)
C
Abdomen bout count
***
*
*
Time (min)
D
Abdomen bout duration (s)
*
*
Time (min)
Wing grooming
E
Wing bout count
Time (min)
F
Wing bout duration (s)
Time (min)

Figure S3

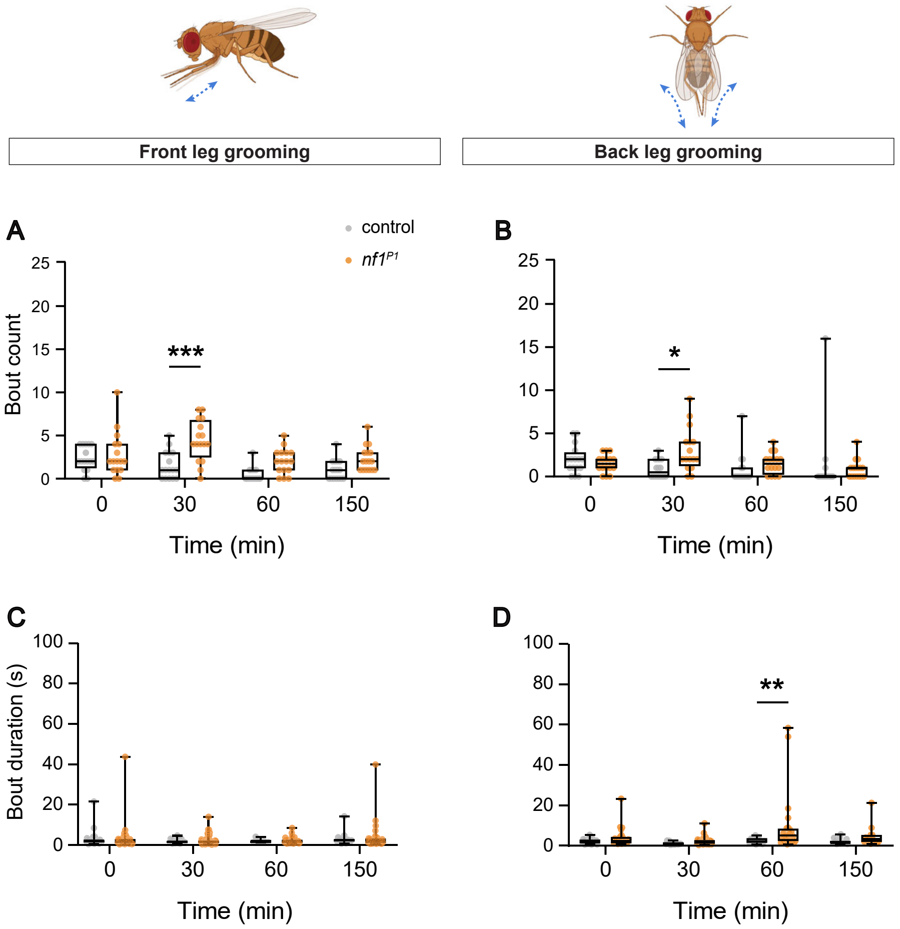

Front leg grooming
Back leg grooming
A
B
C
D
control
nf1P1
Bout count
Bout duration (s)
Time (min)
0
30
60
150
***
*
**


Figure S4

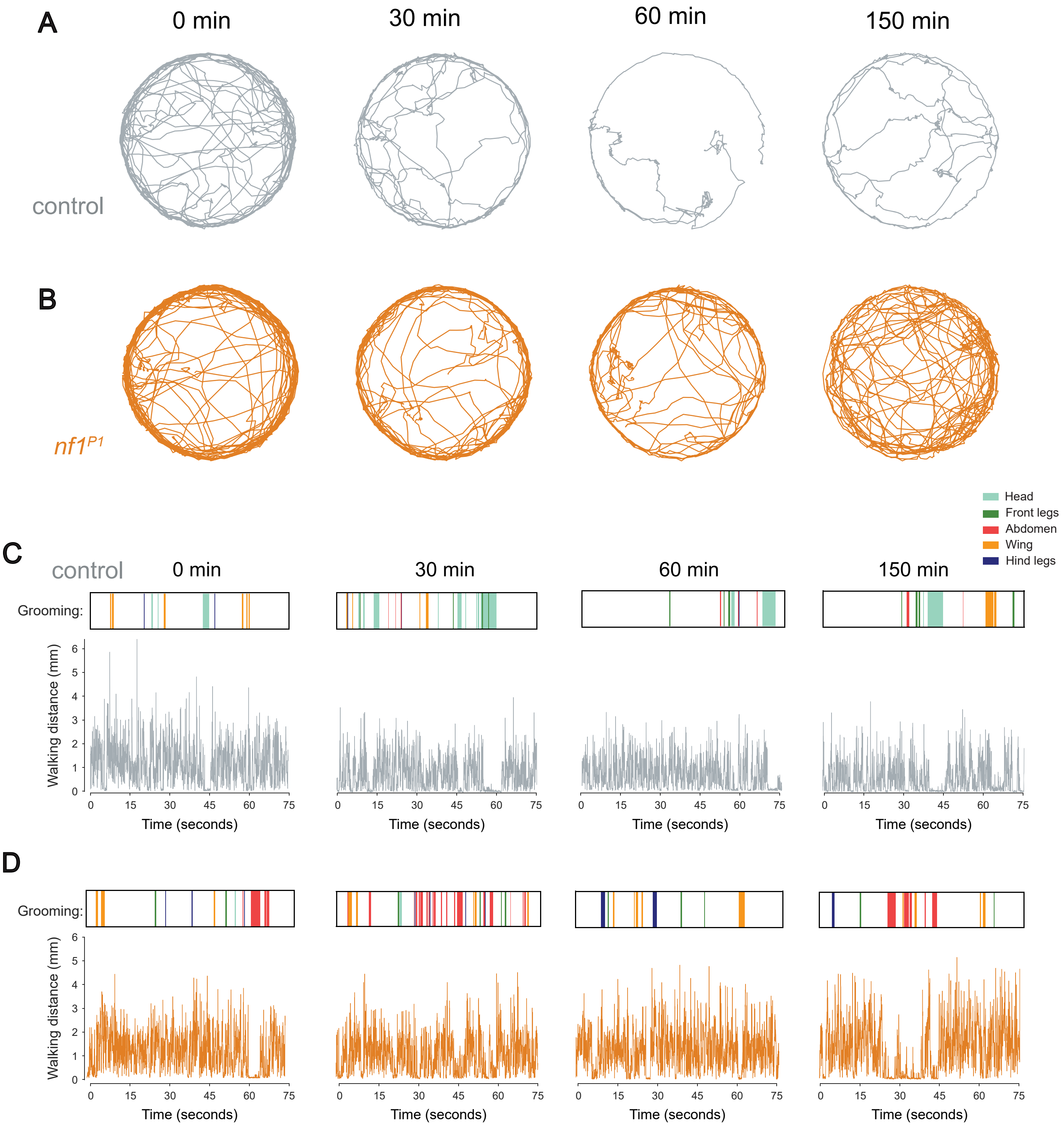

A
0 min
30 min
60 min
150 min
control
B
nf1P1
Head
Front legs
Abdomen
Wing
Hind legs
C
control
0 min
30 min
60 min
150 min
Grooming:
Walking distance (mm)
0
1
2
3
4
5
6
0
15
30
45
60
75
Time (seconds)
D
Grooming:
Walking distance (mm)
Time (seconds)


Figure S5

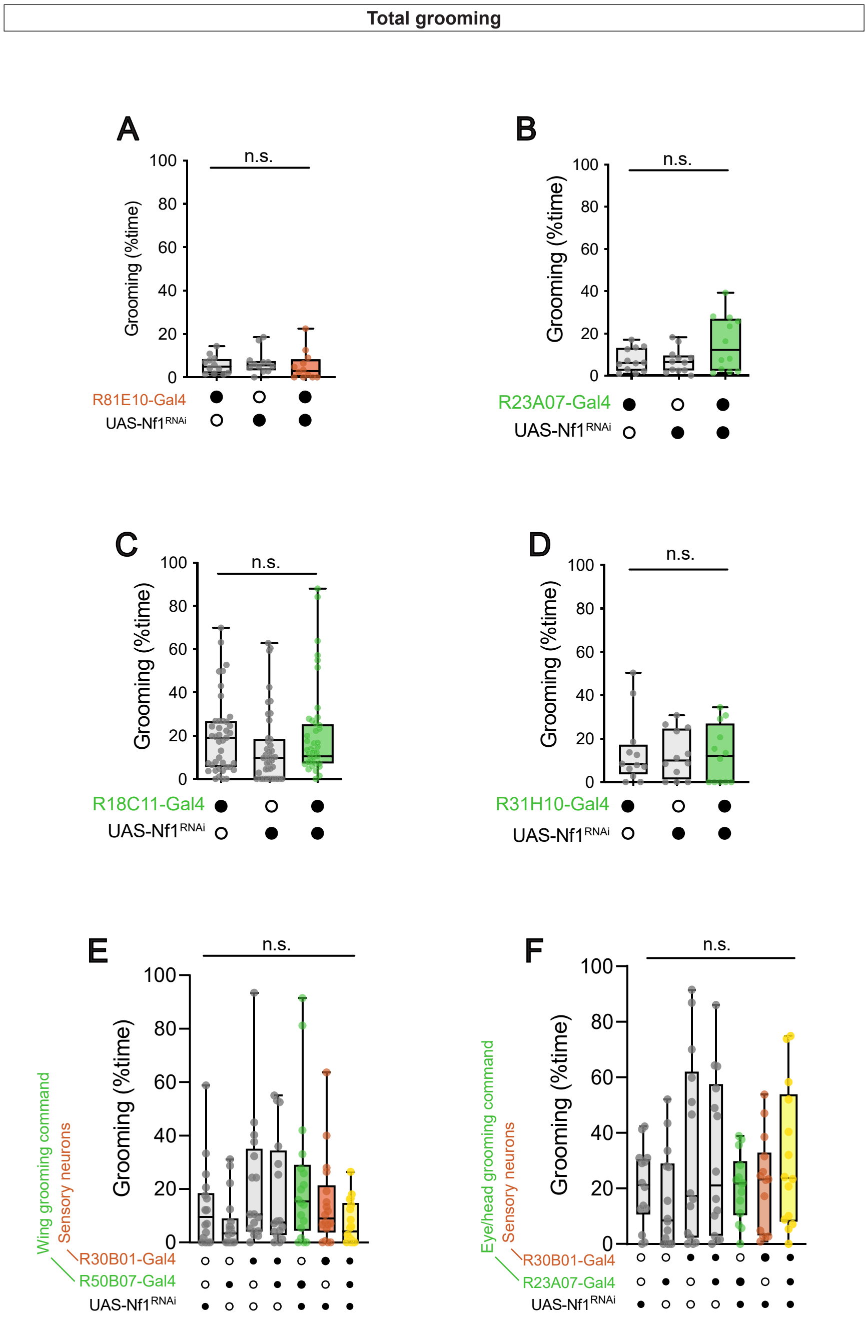
Total grooming
A
B
C
D
E
F
n.s.
Grooming (%time)
R81E10-Gal4
R23A07-Gal4
R18C11-Gal4
R31H10-Gal4
UAS-Nf1RNAi
Wing grooming command
Eye/head grooming command
Sensory neurons
R30B01-Gal4
R50B07-Gal4

Figure S6
